# Antisense transcription of the Neurospora frequency gene is rhythmically regulated by CSP-1 repressor but dispensable for clock function

**DOI:** 10.1101/2022.09.13.507755

**Authors:** Ibrahim A. Cemel, Axel C.R. Diernfellner, Michael Brunner

## Abstract

The circadian clock of *Neurospora crassa* is based on a negative transcriptional/translational feedback loops. The *frequency* (*frq*) gene controls the morning-specific rhythmic transcription of a sense RNA encoding FRQ, the negative element of the core circadian feedback loop. In addition, a long noncoding antisense RNA, *qrf*, is rhythmically transcribed in an evening-specific manner. It has been reported that the *qrf* rhythm relies on transcriptional interference with *frq* transcription and that complete suppression of *qrf* transcription impairs the circadian clock. We show here that *qrf* transcription is dispensable for circadian clock function. Rather, the evening-specific transcriptional rhythm of *qrf* is mediated by the morning-specific repressor CSP-1. Since CSP-1 expression is induced by light and glucose, this suggests a rhythmic coordination of *qrf* transcription with metabolism. However, a possible physiological significance for the circadian clock remains unclear, as suitable assays are not available.

## Introduction

Circadian clocks are widespread biological timing systems that orchestrate and coordinate biochemical pathways, physiology and behavior of organisms in a time-of-day specific manner. They are synchronized by daily recurring cues with the 24 h period of the earth’s rotation and thereby allow the anticipation of changes associated with the geophysical day-night cycle. The core of a circadian clock is based on cell-autonomous transcriptional-translational feedback loops (TTFLs) [1–7].

The circadian clock of the filamentous fungus *Neurospora crassa* is driven by the transcription activator White Collar Complex (WCC), which supports rhythmic transcription of the core clock gene *frequency* (*frq*) and many *clock-controlled genes (ccg’s*) [8, 9]. The intrinsically disordered FRQ protein [10] dimerizes and assembles with FRQ-interacting RNA helicase (FRH) [11–14] and casein kinase 1a (CK1a) [15, 16] forming the FRQ-FRH-CK1a complex, FFC. The FFC inhibits and stabilizes the WCC by facilitating its phosphorylation by CK1a [9, 17]. In the course of a circadian period, FRQ is progressively hyperphosphorylated [18], leading to its inactivation and degradation [19, 20]. The CK1a subunit of the FFC is the major kinase of FRQ [16, 18, 21] but other kinases also contribute to the phosphorylation (for review see: [6, 22]). When the level/activity of FRQ declines, the previously phosphorylated WCC is reactivated by dephosphorylation by protein phosphatases including PP2a [9] and PP5 [23]. The reactivated WCC can now initiate a new circadian cycle of transcription of its target genes. At the *frq* locus it binds about 1.2 kb upstream of the transcription start sites (TSS) to the so-called clock-box (c-box) [24] in order to initiate a new round of *frq* transcription.

The WCC is a heterodimeric transcription factor composed of a WC-1 and a WC-2 subunit [25]. WC-1 contains a LOV-domain photoreceptor that is activated by blue-light [24]. Light-activated WC-1 triggers dimerization of two WCC protomers [26], which then bind to a light response element (LRE) in the core *frq* promoter [24]. The WCC dimer induces high levels of *frq* transcription and thereby synchronizes the circadian clock with light cues [27]. Binding of the light-activated WCC to the LRE induces also transcription of a long noncoding RNA of unknown function that is transcribed towards the c-box [28]. In addition, a promoter located in the 3’ region of the *frq* gene directs transcription of an antisense RNA [28] termed *qrf* [29]. In constant darkness, *qrf* is rhythmically transcribed in antiphase to the overlapping *frq* sense RNA [28, 29]. The *qrf* promoter contains a light response element, qLRE, and the light-activated WCC induces transcription of *qrf* RNA that interferes with and reduces expression of light-induced *frq* sense RNA about 2-fold [29].

The biological function of *qrf* transcription and its impact on the circadian clock are not understood. Kramer et al. [28] replaced the *qrf* promoter at the EcoRV site located 219 bp downstream of the *frq* open reading frame (ORF) (see Figure 1a) by the 3’ UTR of the *clock-controlled gene-2* (*ccg-2*) of *Neurospora*. This manipulation did not affect circadian clock function per se, but was associated with a small (2 h) but significant delay of the phase of the circadian conidiation rhythm. A later analysis by Xue et al. [29] claimed, but did not show, that the EcoRV deletion did not completely abolish *qrf* transcription. They replaced the *qrf* promoter at the BssHII site 143 bp downstream of *frq* ORF with the quinic acid-inducible *qa-2* promoter. When *qa-2*-dependent antisense transcription was either completely repressed or highly induced, the circadian conidiation rhythm of *Neurospora* was abolished. Xue et al. [29] concluded that constitutive antisense transcription at a low level is mechanistically required for the generation of circadian transcriptional rhythms of *frq* sense RNA and therefore essential for the circadian clock. Finally, Li et al. [30] detected a low-amplitude circadian rhythm of *frq* transcription when *qrf* RNA was expressed under control of the induced *qa-2* promoter, suggesting that strongly induced *qrf* transcription did not fully inactivate the circadian clock.

**Figure 1.**
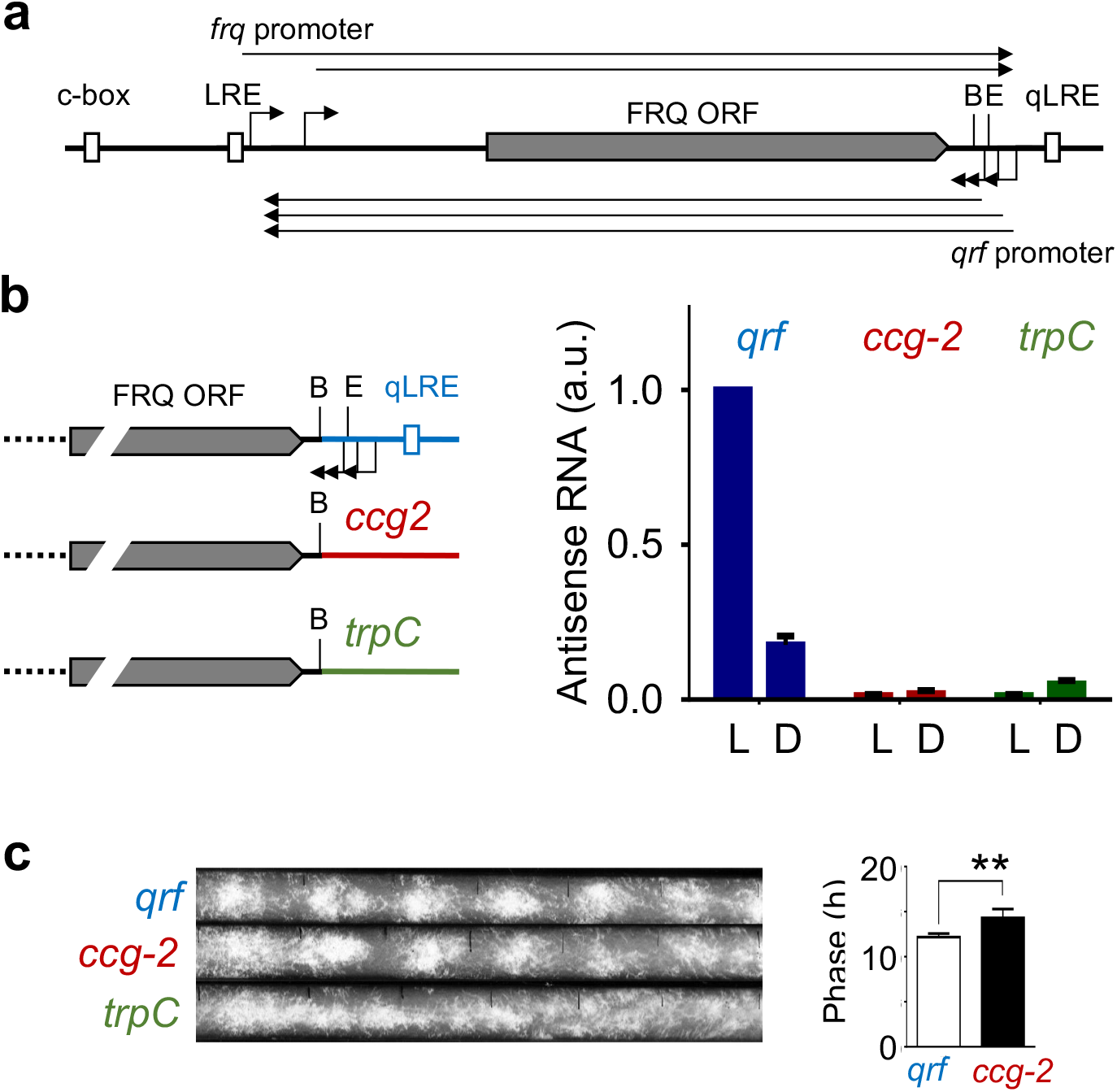
**(a)** Schematic of the *frq* locus. The arrows indicate transcription start sites and the filled box indicates the FRQ ORF. B: BssHII site, E: EcoRV site. **(b)** (left) Schematic representation of the *qrf* mutants. The *WT frq* locus and the *frq* genes with the *ccg-2* and the *trpC* 3’ UTRs, respectively, are depicted. (right) Relative expression levels of *qrf* in the *WT* control and Δ*frqfrq-ccg2 and* Δ*frq,frq-trpC* strains. The samples were harvested in constant light (LL) and 22 h after a light to dark transfer (DD22). *Actin* was used as an internal reference. Error bars represent ±SEM (n=3). **(c)** Representative race tube assays of the indicated strains. frq^wt^: τ = 22.81 ± 0.19 h, Δ*frq,frq-ccg2:* τ = 22.39 ± 0.70 h, Δ*frqfrq-trpC:* arrhythmic (n=6). τ: period length. The bar plot shows the phases of the *WT* control and Δ*frq,frq-ccg2*. Error bars represent ±SEM (n=4-6). ** indicates p-value <0.01.

We show here that the *qrf* promoter dispensable for circadian rhythmicity. Rather, the circadian clock is highly sensitive to changes in the 3’-UTR of *frq* RNA that affect RNA turnover. We also show that the glucose and light-induced transcriptional repressor CSP-1 supports rhythmic expression of *qrf* in antiphase to *frq*.

## Materials and Methods

### Neurospora strains and culture conditions

*Neurospora* wild-type strain was acquired from Fungal Genetics Stock Center (FGSC) with *bd* stock number #2489. For race tube assay, strains with *ras-1^bd^* mutation were used [31]. Since the conventional *frq* deletion strain *frq^10^* [32] contains the enhancer site c-box and the *qrf* LRE, we created a full knock-out of the *frq* locus in this study. The Δ*frq bd, his-3* strain was created with the yeast *in vivo* recombination system as described previously [33] using the following knockout cassette primers:

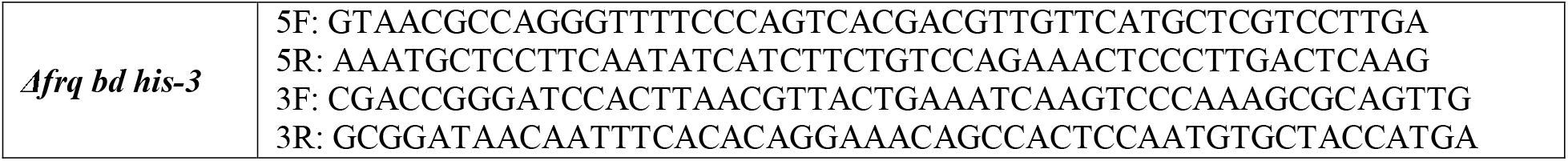

The plasmids for the *qrf* knockout strains were created with the primers:

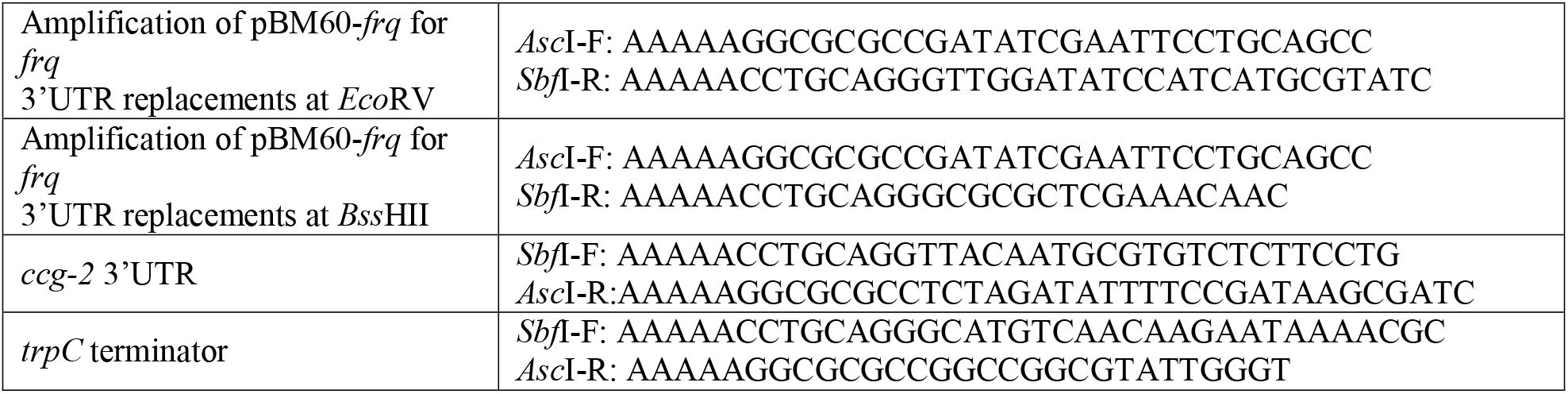

The Δ*frq bd, his-3* strain was used for transformation. Briefly, 5-7 days old conidia were harvested and washed and pelleted at 2600g at 4°C for 10 min with 50 ml of 1 M cold sorbitol. The conidial pellet was mixed with 1-2 μg linearized plasmid DNA and incubated on ice for 10 min. Electroporation was applied at 1.5 kV/cm, 25 μF, 600 Ω. The cells were immediately resuspended in 1 ml of 1 M cold sorbitol and plated onto Vogel’s solid media (1x Vogel’s, 1% (w/v) agar, 1x FGS (20% (w/v) sorbose, 0.5% (w/v) glucose, 0.5% (w/v) fructose). After incubation at 30°C for 3-5 days, the single colonies were picked. *N. crassa* cultures were grown in standard growth medium (2% glucose, 0.5% L-arginine, 1x Vogel’s medium, and 10 ng/mL biotin) at 25°C with shaking at 115 rpm. In light-induction experiments, the strains were grown into mats in petri plates with 20 ml of standard medium. Mycelial discs (1 cm) were cut out and grown in standard growth medium for 1 day.

### RNA preparation and cDNA synthesis

RNA was isolated with peqGOLD TriFAST (PeqLab). The reverse transcription was carried out with QuantiTect Reverse Transcription Kit (Qiagen) with indicated primers (Table 3) following manufacturer’s instructions. Relative transcript levels were quantified by quantitative real-time PCR in 96-well plates with LightCycler 480 (Roche). The reaction was set by using qPCRBIO Probe Mix Hi-ROX (Nippongenetics) and TaqMan (5’: 6-FAM, 3’: TAMRA) or UPL (Universal Probe Library, Roche) probes. Primers and probes are listed below. Three replicates were used to calculate the mean threshold cycle (Ct) value. The relative enrichments were quantified relative to a housekeeping gene using the reverse transcription primers for cDNA synthesis:

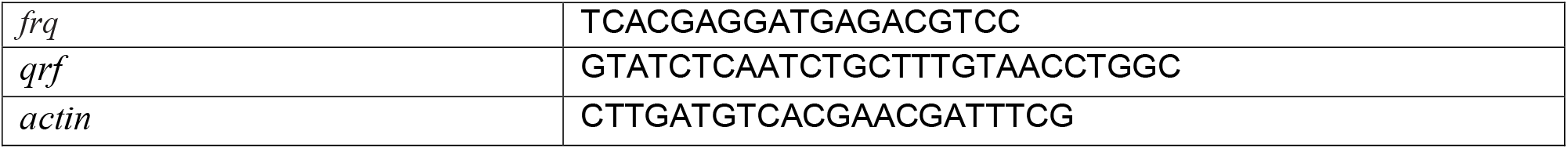

And primers for qPCR:

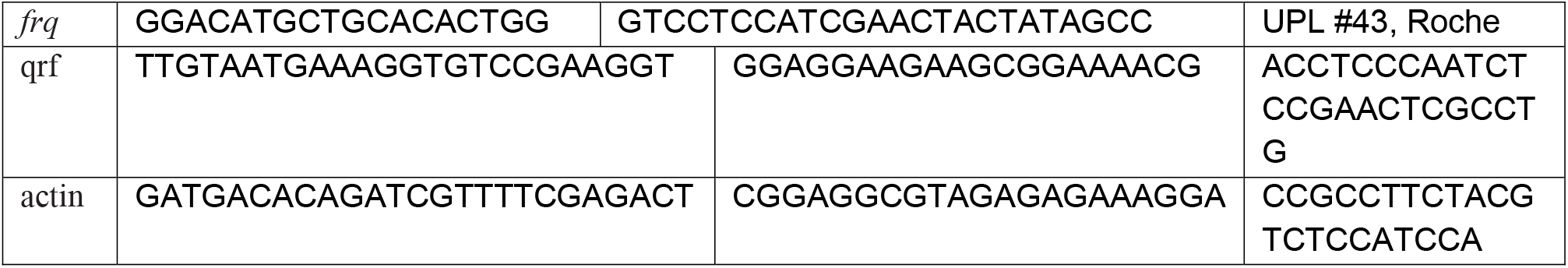

### Race-tube assay

The conidial suspension was inoculated from one end of the autoclaved glass tubes containing the media (1x Vogel’s, 0.1% glucose (w/v), 0.17% arginine (w/v), 50 ng/ml biotin and 2% agar (w/v)) and was grown for 1 day prior to light to dark transfer. The conidial growth fronts were marked every 24 h.

### Live-cell bioluminescence monitoring

Sorbose medium (1x FGS (0.05% fructose (w/v), 0.05% glucose (w/v), 1% sorbose (w/v)), 1x Vogel’s, 1% agarose (w/v), 10 ng/ml biotin and 75 μM firefly luciferin) was used for the livecell bioluminescence assay. 96-well plates were inoculated with 3×10^4^ conidia per well and incubated in dark at 25°C for 2 days. The bioluminescence signal was recorded in constant darkness or in light-dark cycles at 25°C for the indicated time windows with a multilabel plate reader. For dark recordings, the cells were synchronized by 1 h light pulse at 100 μmol photons m^-2^ s^-1^ prior to the measurement in constant darkness. 0.3% glucose as carbon source was used when high-glucose conditions are indicated. The light intensity titration (Figure 3e) was performed as described [34].

## Results and discussion

### *Antisense transcription at* frq *locus is not required for rhythmicity*

Previous studies [28, 29, 35] and RNA-Seq data [36] indicate that the *frq* locus directs expression of multiple species of overlapping sense and antisense transcripts (Figure 1a). Light-induced expression of *frq* and its antisense transcript, *qrf*, are driven from their corresponding light response elements, LRE and qLRE, respectively [24, 29, 37, 38]. The majority of sense *frq* RNA species terminate between the qLRE and the annotated transcription start sites (TSSs) of the *qrf* promoter [39], and *qrf* transcripts terminate within the region of annotated TSSs of the *frq* promoter (Figure 1a).

To analyze the impact of antisense transcription on the circadian clock we replaced the *qrf* promoter with two different transcription termination sequences, the 3’ UTR of the *clock-controlled gene-2 (ccg-2*) of *Neurospora crassa* [28] and the *trpC* terminator of *Aspergillus nidulans* [40], respectively. The sequences were inserted at the BssHII site to ensure deletion of all mapped *qrf* TSSs (Figure 1b, left). The modified *frq* genes and an unmodified *frq* gene with its natural *qrf* sequence (*frq-qrf* control), respectively, were inserted into the *Neurospora* genome at the *his-3* locus of a Δ*frq* strain (see Material and Methods). RNA directed from these genes was quantified by qRT-PCR (Figure 1b, right). In the *frq-qrf* control strain, antisense RNA was expressed at a high level in light and at a ~5-fold lower level in the dark. In contrast, in the *frq-ccg-2* and *frq-trpC* strains the antisense transcript levels were significantly reduced, indicating that deletion of the *qrf* promoter at the BssHII site compromised the expression of antisense RNA in light and in dark.

We then assessed the conidiation rhythms of the *frq-qrf* control strain and the mutant strains lacking antisense transcription. The *frq-qrf* control strain and *frq-ccg-2* strain exhibited rhythmic conidiation in constant darkness (DD) but the phase of conidiation of the *frq-ccg-2* strain was delayed by 2 h (Figure 1c). The observed phase delay is consistent with a previous report [28], in which the *qrf* promoter had been replaced at the EcoRV site by the 3’-UTR of *ccg-2*. In contrast, conidiation in DD was arrhythmic in the *frq-trpC* strain, indicating that the circadian clock was severely compromised. Thus, despite the *frq-ccg-2* and *frq-trpC* strains being both deficient in antisense transcription, *frq-ccg-2* was rhythmic while *frq-trpC* was apparently arrhythmic.

Since the *frq* mRNA terminates downstream of the mapped *qrf* TSSs [39], replacement of the *qrf* promoter by *ccg-2* or *trpC* altered the 3’ UTR of the *frq* sense transcript. We therefore asked how the choice of the DNA sequence, which was used to replace the *qrf* promoter, affected the stability of *frq* mRNA. To assess RNA turnover under physiological conditions, we allowed accumulation of high levels of the *frq* RNA by growing mycelial cultures in constant light. The light-induced transcription of *frq* was then shut-down by a transfer of the cultures to the dark and the decrease of *frq* RNA was then measured over a time course of 120 min (Figure 2a). The *frq* RNA levels in the *frq-qrf* control strain decreased with a half-time (t_1/2_) of 15 min, confirming that *frq* RNA is unstable [41]. The level of the *frq-ccg-2* transcript decreased with t_1/2_ ~ 25 min while the *frq-trpC* transcript was substantially more stable (t_1/2_ ~ 1 h). The data demonstrate, not surprisingly, that the 3’ UTRs affect the stability and hence the expression level of *frq* mRNA.

**Figure 2.**
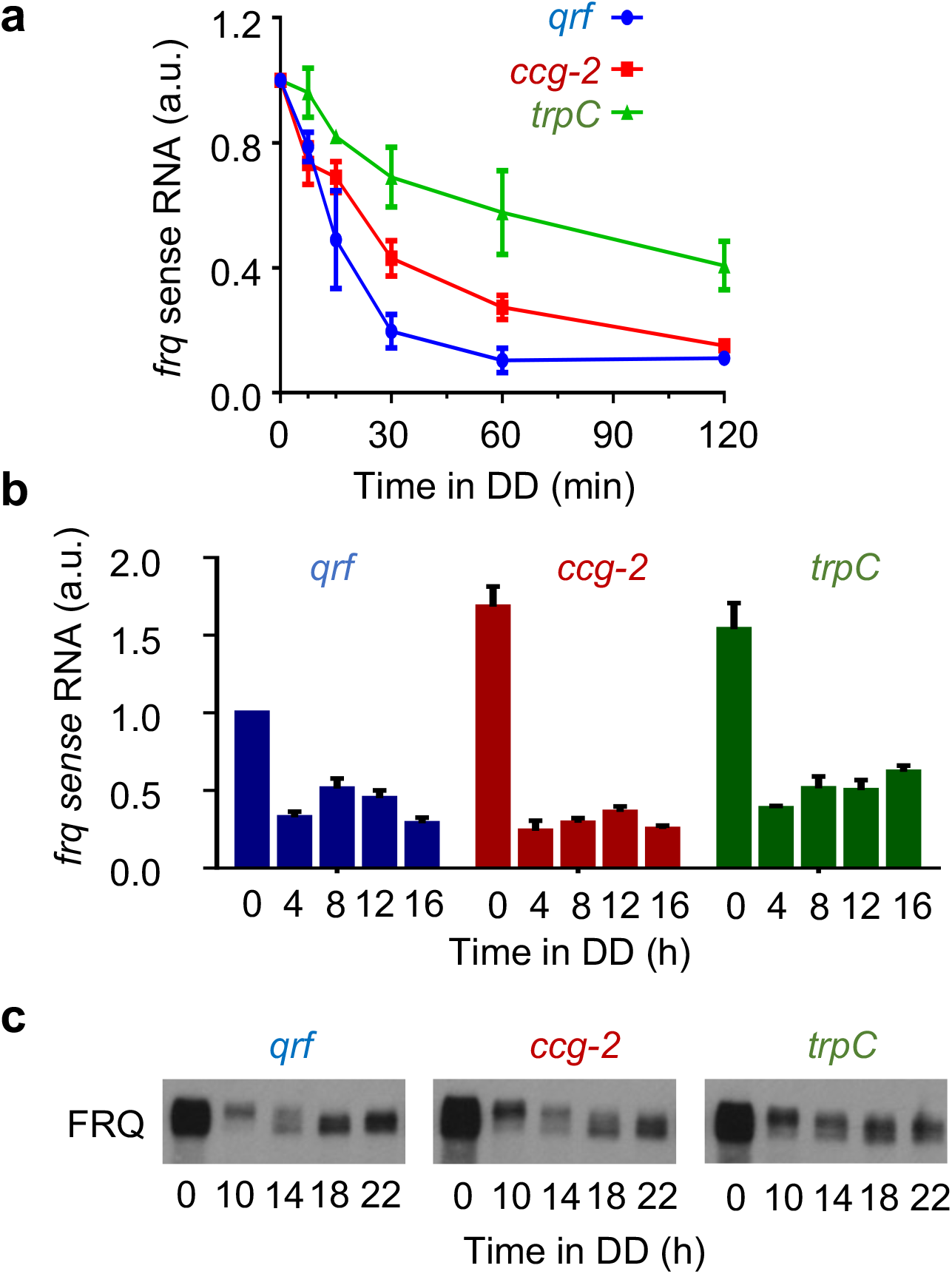
**(a)** Stability of *frq* mRNA in *WT* and *qrf* deficient strains. Strand-specific RT-qPCR results showing the frq mRNA levels in light (0 h) and in darkness at indicated time points (min). The transcript levels were normalized to the respective light value, *actin* was used as an internal reference. Error bars indicate ±SEM (n=3). **(b)** Rhythmic expression of *frq* in *wt* and *qrf* deficient strains. Strand-specific RT-qPCR results showing the frq mRNA levels in light (0 h) and in darkness at indicated time points (h). Relative *frq* levels were normalized to the *wt* levels in light (0 h). Error bars indicate ±SEM (n=3). **(c)** Western blot analysis of FRQ expression profiles in the Δfrq,frq-ccg2 and Δfrq,frq-trpC strains in DD at the indicated time points (n=3).

We then measured *frq* RNA and FRQ protein in light and after LD transfer of mycelial cultures (Figure 2b). In light (t = 0 h), *frq* transcript levels in the *frq-ccg-2* and *frq-trpC* strains accumulated at a higher level than in the *frq-qrf* control strain, confirming previous reports that replacement of the antisense promoter results in accumulation of elevated levels of sense RNA [29]. In constant darkness, the *frq-qrf* control strain expressed *frq* sense RNA in circadian fashion, with a peak about 12 h after transfer of the mycelial cultures into the dark (Figure 2b). FRQ protein displayed circadian abundance and phosphorylation rhythms (Figure 2c). In the *frq-ccg-2* strain, *frq* RNA and FRQ protein also displayed circadian rhythms in DD, however, their circadian phase was delayed (Figure 2b, c). In the *frq-trpC* strain, the levels of *frq* and FRQ in dark were elevated and did not display apparent rhythms on the time frame of the experiment (Figure 2b, c). Hence, the delayed conidiation rhythm of *frq-ccg-2* and the arrhythmic conidiation of *frq-trpC* strains correlate with the expression profiles of *frq* and FRQ in these strains after LD transfer.

Together, our results show that antisense (*qrf*) transcription at the *frq* locus is not required for circadian rhythmicity. Rather, the circadian clock is sensitive to changes in the 3’ UTR that stabilize *frq* RNA, emphasizing that rapid turnover of *frq* RNAis crucial for clock function.

### *Transcription dynamics of* qrf *in dark and in response to light*

Next, we characterized the transcriptional properties of the *qrf* promoter using luciferase reporter constructs. Three TSSs of the *qrf* promoter were previously mapped, two between the BssHII and EcoRV sites and one immediately downstream of the EcoRV site (Figure 3a). To include all TSSs, we fused the *qrf* promoter immediately after the stop codon of the FRQ open reading frame to *luc-PEST* (Figure 3a), encoding a destabilized luciferase gene [42, 43].The DNA sequence transcribed by the *qrf* promoter contains several ATG codons, and 3 of these are present in the 5’-UTR of the chimeric *qrf-lucPEST* transcription unit. We changed these 3 upstream ATGs in the reporter gene to CTGs to ensure optimal translation of *lucPEST* ORF. The modified *qrf-lucPEST* reporter supported efficient bioluminescence expression and exhibited a low-amplitude circadian rhythm (Figure 3b). We also generated *lucPEST* reporter constructs with *qrf* promoters truncated at the EcoRV and BssHII site, respectively (Figure 3a). In comparison to the transcriptional activity of the full-length *qrf-lucPEST* reporter, the bioluminescence levels supported by *qrf* promoters truncated at EcoRV and BssHII, respectively, were substantially reduced (Figure 3b). The residual activity was arrhythmic and similarly low for the EcoRV and BssHII truncations. The data suggest that truncations at either restriction site abolishes *qrf* rhythms and transcription levels to the same extent.

**Figure 3.**
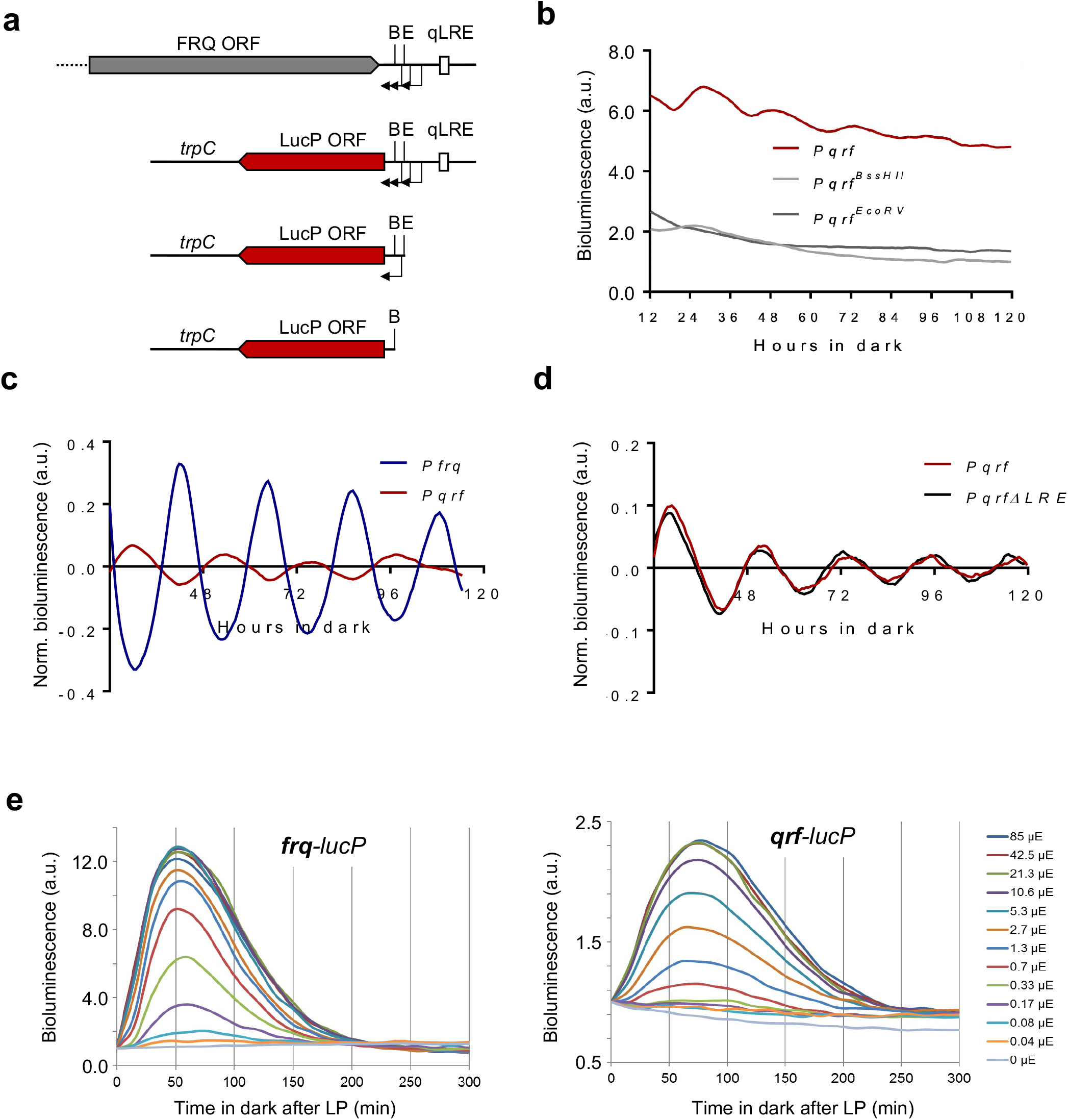
**(a)** Schematic representation of the *luciferase* reporter constructs together with the *frq* locus. The arrows indicate transcription start sites. The filled boxes represent FRQ ORF (grey) or LUC ORF (red). B: BssHII site, E: EcoRV site. **(b)** Representative normalized bioluminescence recordings of luciferase reporters driven by the complete qrf promoter, truncated qrf promoters at EcoRV or BssHII restriction sites. The reporters were expressed in wt bd. Two clones and their means are depicted. **(c)** Rhythmic *qrf* promoter activity. Normalized luciferase activity of the *qrf* promoter in comparison to the *frq* promoter activity. The bioluminescence activities were recorded in constant darkness after synchronization via light to dark transfer. **(d)** Normalized *qrf* and *qrfΔLRE* driven luciferase activities and in constant darkness. **(e)** Differential saturation of *frq* and *qrf* promoters. Strains expressing *frq-lucPEST* and *qrf-lucPEST* reporter genes were exposed to a 1-min light pulse of the indicated intensities. Luciferase activity at light pulse was used for normalization (n=3).

We then compared the transcription rhythms of the isolated *qrf* (antisense) reporter with the rhythm supported by a *frq* (sense) reporter gene (Figure 3c). The expression levels of the *qrf-lucPEST* reporter oscillated in antiphase to the circadian rhythm of the *frq-lucPEST* reporter.

Light-dependent expression of *qrf* is controlled by the qLRE [29, 38]. To assess whether the qLRE is required for the anti-phasic transcription rhythm in the dark we mutated the sequence in the *qrf-lucPEST* reporter and measured luciferase expression in the corresponding reporter strain, *qrfΔqLRE*. The qLRE mutation did not affect the evening-specific circadian expression rhythm of the *qrfΔqLRE* promoter (Figure 3d).

Finally, to compare the responsiveness and sensitivity of the *frq* and *qrf* promoters to light, the *frq-lucPEST* and *qrf-lucPEST* reporter strains were exposed to 1 min light pulses (LPs) of different intensity and bioluminescence was then recorded for 300 min (Figure 3e). The *frq-lucPEST* reporter reached peak expression levels about 50 min after the LP, while the *qrf-lucPEST* reporter reached maximal expression levels after about 75 min. Furthermore, expression levels of the *frq-lucPEST* reporter saturated at LP intensities above 2.7 μmol photon s^-1^ m^-2^ while the *qrf* promoter saturated at much higher LP intensities (>21.3 μmol photon s^-1^ m^-2^). The data indicate that the *qrf* promoter is less responsive and less sensitive to light cues.

### *The* qrf *promoter is controlled by CSP-1*

In constant darkness, clock-controlled genes of *Neurospora* are rhythmically expressed in mainly two phases, subjective morning and evening, respectively [44, 45]. Morning-specific genes are expressed approximately in phase with the activity profile of the transcription activator WCC, while many evening-phased genes are regulated by the transcription repressor CSP-1 [45]. Since the transcriptional activity of *qrf* is evening-specific, i.e. in antiphase to the rhythm of the WCC-driven *frq* promoter, we asked whether the *qrf* promoter is controlled by the morning-specific CSP-1 repressor. Analysis of CSP-1 ChIP-Seq [46] and WC-2 ChIP-Seq [47] datasets revealed that CSP-1 binds overlapping with the LRE and qLRE of the *frq* and *qrf* promoters, respectively, and also close to the clock-box (c-box) of *frq* (Figure 4a). We have previously shown that *frq* expression is elevated and phase-delayed in a Δ*csp-1* strain [46]. To assess the impact of CSP-1 on the *qrf* promoter we analyzed expression of *qrf-lucPEST* in a Δ*csp-1* strain. Expression of *qrf-lucPEST* was elevated (*P* < 0.05) and its circadian rhythm was abolished. The data suggest that *qrf* is constitutively activated by at least one unknown transcription factor (TF). The activity of this TF is antagonized by CSP-1, which represses *qrf* transcription. Since CSP-1 is rhythmically expressed with a morning-specific phase, it generates an anti-phasic evening-specific *qrf* rhythm (Figure 2d). Together these results demonstrate that the isolated *qrf* promoter displays an evening-specific expression rhythm, which is dependent on the morning-specific activity of the repressor CSP-1.

**Figure 4.**
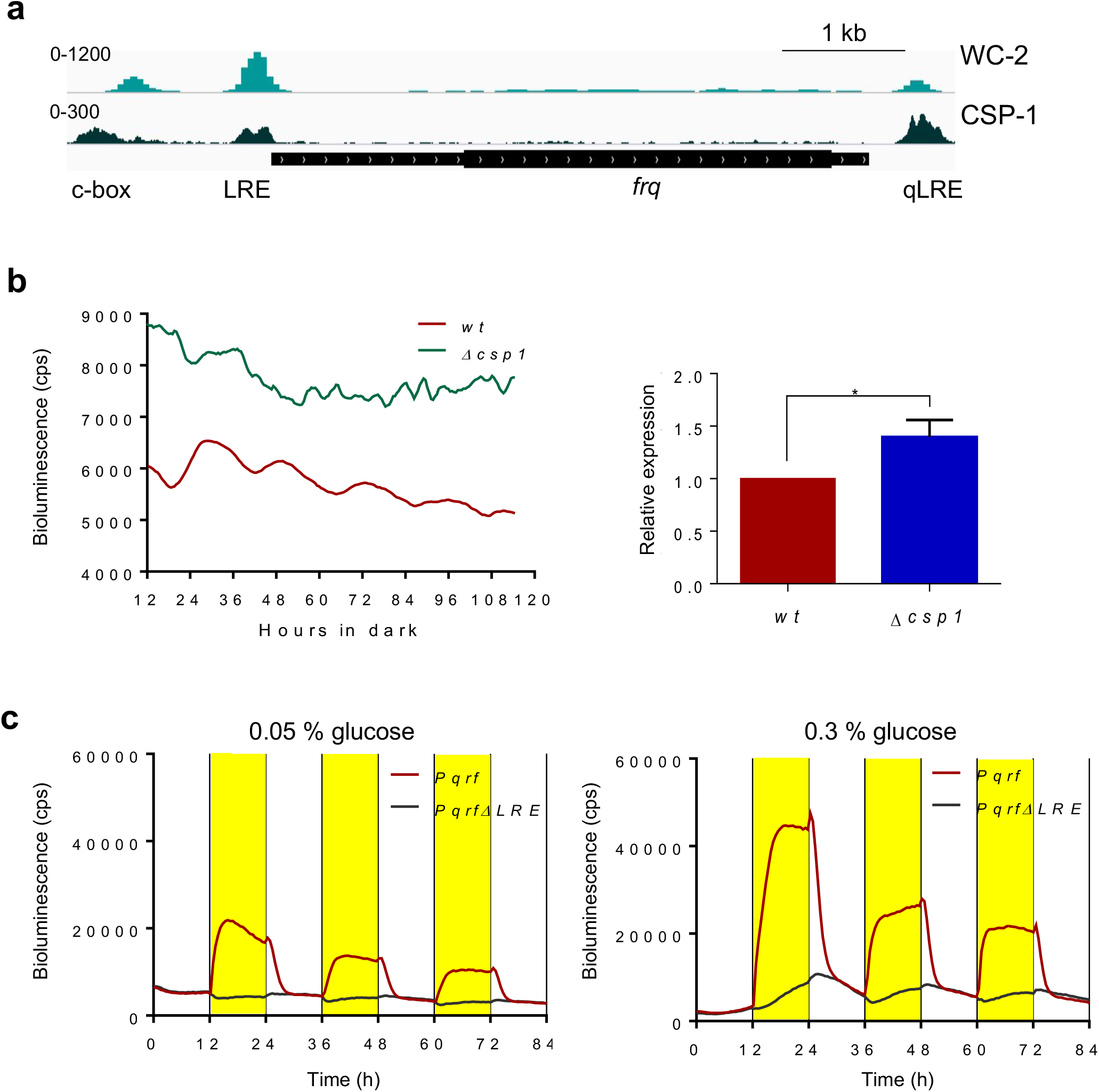
**(a)** WC-2 and CSP-1binding at the *frq* locus. ChIP-Seq datasets were published in [46] and [47], respectively. **(b)** Luciferase activity of *qrf* promoter in *WT* and Δ*csp-1* strains. Following light to dark transfer, the bioluminescence activities were recorded in constant darkness. The quantification of three independent experiments are shown. Error bars represent ±SEM (n=3). * indicates p-value <0.05. **(c)** Representative bioluminescence measurement of the *qrf-lucPEST* and *qrfΔLRE-lucPEST* reporter in low (0.05%) and high (3%) glucose concentrations in 12 h dark (white) −12 h light (yellow) (LD) cycles.

CSP-1 is a short-lived repressor that is expressed in light-dependent fashion under transcriptional control of the WCC, and independently of WCC also in glucose-dependent fashion [48]. To characterize the impact of CSP-1 on *qrf* transcription we examined *qrf-lucPEST* and *qrf*Δ*qLRE-lucPEST* reporter strains. The strains were cultured in 96-well plates on agar medium with low and high glucose concentration. The cultures were exposed to repetitive 12h:12h LD cycles and bioluminescence profiles were recorded for 84 h (Figure 4a, b). The *qrf-luc-PEST* strain displayed light-driven luciferase activity (bioluminescence) at low and high glucose concentration. On low glucose medium (Figure 3a, left) the bioluminescence levels decreased rapidly after the LD transition and approached dark levels within ~4 h. On high glucose medium (Figure 3b, right), the decrease in bioluminescence after the LD transition was bi-phasic: the initial rapid decrease of bioluminescence within the first 4 h in the dark was followed by a slower decrease during the 4 −12 h time period after the LD-transition. To our surprise, the bioluminescence supported by *qrf*Δ*qLRE-lucPEST* was modulated by light in a glucose-dependent manner (Figure 4c). At low glucose concentration *qrf*Δ*qLRE-lucPEST* expression was essentially unaffected by light (Figure 4c, left). In contrast, at high glucose concentration in the medium, *qrf*Δ*qLRE-lucPEST* expression displayed a specific response to light despite the absence of a functional qLRE (Figure 4c, right): after about 1 h in light, the bioluminescence supported by *qrf*Δ*qLRE-lucPEST* increased steadily. After LD-transition, the bioluminescence kept increasing for about 2 h and then declined steadily, resulting in a sawtooth-like bioluminescence rhythm that was delayed relative to the DL- and LD-transitions. Interestingly, the decrease in the dark of *qrf*Δ*qLRE-lucPEST* expression superimposed with the slow decrease of *qrf-lucPEST* activity observed 4 - 8 h after LD-transition (Figure 4c). The data suggest that light supports the expression of an unknown, glucose-dependent transcription activator of *qrf*, which accumulates steadily during the light phase and is degraded during the dark phase. Hence, the *qrf* promoter is regulated in complex manner by at least two environmental cues, light and glucose. Light-dependent expression is directly controlled by the WCC via the qLRE while glucose-dependent expression is controlled by an unknown activator and by the rhythmically expressed repressor CSP-1. Activity and presumably expression levels of both, activator and repressor, are dependent on light and glucose.

Taken together, our result demonstrate that antisense transcription is not required for the function of the circadian clock in constant darkness (Figure 1 and 2). Rather, the impact *qrf* transcription on *frq* transcription in light via WCC [28, 29] (Figure 3), the direct impact of glucose via CSP1 on *frq* [46] and *qrf* transcription (Figure 4a, b), and the indirect impact of glucose and light on *qrf* via an unknown transcription activator (Figure 4c) suggests that antisense transcription may help finetune and coordinate the light-dark phase of *frq* expression with the metabolic state of the fungus.

## Acknowledgements

This work was supported by the Deutsche Forschungsgemeinschaft (collaborative research center TRR186).

## Conflict of Interest Statement

The authors have no potential conflicts of interest with respect to the research, authorship, and/or publication of this article.

## Notes

### Competing Interest Statement

The authors have declared no competing interest.

## References

1. Dunlap, J. C. & Loros, J. J. (2017) Making Time: Conservation of Biological Clocks from Fungi to Animals, Microbiol Spectr. 5.

2. Rosbash, M. (2009) The implications of multiple circadian clock origins, PLoS Biol. 7, e62.

3. Hogenesch, J. B. & Ueda, H. R. (2011) Understanding systems-level properties: timely stories from the study of clocks, Nat Rev Genet. 12, 407–16.

4. Takahashi, J. S. (2017) Transcriptional architecture of the mammalian circadian clock, Nat Rev Genet. 18, 164–179.

5. Gallego, M. & Virshup, D. M. (2007) Post-translational modifications regulate the ticking of the circadian clock, Nat Rev Mol Cell Biol. 8, 139–48.

6. Diernfellner, A. C. R. & Brunner, M. (2020) Phosphorylation Timers in the Neurospora crassa Circadian Clock, J Mol Biol. 432, 3449–3465.

7. Partch, C. & Brunner, M. (2022) How circadian clocks keep time: the discovery of slowness, FEBS Lett. 596, 1613–1614.

8. Dunlap, J. C. (1999) Molecular bases for circadian clocks, Cell. 96, 271–90.

9. Schafmeier, T., Haase, A., Kaldi, K., Scholz, J., Fuchs, M. & Brunner, M. (2005) Transcriptional feedback of Neurospora circadian clock gene by phosphorylation-dependent inactivation of its transcription factor, Cell. 122, 235–46.

10. Pelham, J. F., Dunlap, J. C. & Hurley, J. M. (2020) Intrinsic disorder is an essential characteristic of components in the conserved circadian circuit, Cell Commun Signal. 18, 181.

11. Cheng, P., He, Q., He, Q., Wang, L. & Liu, Y. (2005) Regulation of the Neurospora circadian clock by an RNA helicase, Genes Dev. 19, 234–41.

12. Hurley, J. M., Larrondo, L. F., Loros, J. J. & Dunlap, J. C. (2013) Conserved RNA helicase FRH acts nonenzymatically to support the intrinsically disordered neurospora clock protein FRQ, Mol Cell. 52, 832–43.

13. Lauinger, L., Diernfellner, A., Falk, S. & Brunner, M. (2014) The RNA helicase FRH is an ATP-dependent regulator of CK1a in the circadian clock of Neurospora crassa, Nat Commun. 5, 3598.

14. Cheng, P., Yang, Y., Heintzen, C. & Liu, Y. (2001) Coiled-coil domain-mediated FRQ-FRQ interaction is essential for its circadian clock function in Neurospora, EMBO J. 20, 101–8.

15. Gorl, M., Merrow, M., Huttner, B., Johnson, J., Roenneberg, T. & Brunner, M. (2001) A PEST-like element in FREQUENCY determines the length of the circadian period in Neurospora crassa, EMBO J. 20, 7074–84.

16. Marzoll, D., Serrano, F. E., Shostak, A., Schunke, C., Diernfellner, A. C. R. & Brunner, M. (2022) Casein kinase 1 and disordered clock proteins form functionally equivalent, phospho-based circadian modules in fungi and mammals, Proc Natl Acad Sci U S A. 119.

17. Schafmeier, T., Kaldi, K., Diernfellner, A., Mohr, C. & Brunner, M. (2006) Phosphorylation-dependent maturation of Neurospora circadian clock protein from a nuclear repressor toward a cytoplasmic activator, Genes Dev. 20, 297–306.

18. Querfurth, C., Diernfellner, A. C., Gin, E., Malzahn, E., Hofer, T. & Brunner, M. (2011) Circadian conformational change of the Neurospora clock protein FREQUENCY triggered by clustered hyperphosphorylation of a basic domain, Mol Cell. 43, 713–22.

19. He, Q., Cheng, P., Yang, Y., He, Q., Yu, H. & Liu, Y. (2003) FWD1-mediated degradation of FREQUENCY in Neurospora establishes a conserved mechanism for circadian clock regulation, EMBO J. 22, 4421–30.

20. Larrondo, L. F., Olivares-Yanez, C., Baker, C. L., Loros, J. J. & Dunlap, J. C. (2015) Circadian rhythms. Decoupling circadian clock protein turnover from circadian period determination, Science. 347, 1257277.

21. Querfurth, C., Diernfellner, A., Heise, F., Lauinger, L., Neiss, A., Tataroglu, O., Brunner, M. & Schafmeier, T. (2007) Posttranslational regulation of Neurospora circadian clock by CK1a-dependent phosphorylation, Cold Spring Harb Symp Quant Biol. 72, 177–83.

22. Diernfellner, A. C. & Schafmeier, T. (2011) Phosphorylations: Making the Neurosporacrassa circadian clock tick, FEBS Lett. 585, 1461–6.

23. Cha, J., Chang, S. S., Huang, G., Cheng, P. & Liu, Y. (2008) Control of WHITE COLLAR localization by phosphorylation is a critical step in the circadian negative feedback process, EMBO J. 27, 3246–55.

24. Froehlich, A. C., Liu, Y., Loros, J. J. & Dunlap, J. C. (2002) White Collar-1, a circadian blue light photoreceptor, binding to the frequency promoter, Science. 297, 815–9.

25. Cheng, P., Yang, Y., Gardner, K. H. & Liu, Y. (2002) PAS domain-mediated WC-1/WC-2 interaction is essential for maintaining the steady-state level of WC-1 and the function of both proteins in circadian clock and light responses of Neurospora, Mol Cell Biol. 22, 517–24.

26. Malzahn, E., Ciprianidis, S., Kaldi, K., Schafmeier, T. & Brunner, M. (2010) Photoadaptation in Neurospora by competitive interaction of activating and inhibitory LOV domains, Cell. 142, 762–72.

27. Linden, H. & Macino, G. (1997) White collar 2, a partner in blue-light signal transduction, controlling expression of light-regulated genes in Neurospora crassa, EMBO J. 16, 98–109.

28. Kramer, C., Loros, J. J., Dunlap, J. C. & Crosthwaite, S. K. (2003) Role for antisense RNA in regulating circadian clock function in Neurospora crassa, Nature. 421, 948–52.

29. Xue, Z., Ye, Q., Anson, S. R., Yang, J., Xiao, G., Kowbel, D., Glass, N. L., Crosthwaite, S. K. & Liu, Y. (2014) Transcriptional interference by antisense RNA is required for circadian clock function, Nature. 514, 650–3.

30. Li, N., Joska, T. M., Ruesch, C. E., Coster, S. J. & Belden, W. J. (2015) The frequency natural antisense transcript first promotes, then represses, frequency gene expression via facultative heterochromatin, Proc Natl Acad Sci U S A. 112, 4357–62.

31. Belden, W. J., Larrondo, L. F., Froehlich, A. C., Shi, M., Chen, C. H., Loros, J. J. & Dunlap, J. C. (2007) The band mutation in Neurospora crassa is a dominant allele of ras-1 implicating RAS signaling in circadian output, Genes Dev. 21, 1494–505.

32. Aronson, B. D., Johnson, K. A., Loros, J. J. & Dunlap, J. C. (1994) Negative feedback defining a circadian clock: autoregulation of the clock gene frequency, Science. 263, 1578–84.

33. Colot, H. V., Park, G., Turner, G. E., Ringelberg, C., Crew, C. M., Litvinkova, L., Weiss, R. L., Borkovich, K. A. & Dunlap, J. C. (2006) A high-throughput gene knockout procedure for Neurospora reveals functions for multiple transcription factors, Proc Natl Acad Sci U S A. 103, 10352–10357.

34. Cesbron, F., Oehler, M., Ha, N., Sancar, G. & Brunner, M. (2015) Transcriptional refractoriness is dependent on core promoter architecture, Nat Commun. 6, 6753.

35. Colot, H. V., Loros, J. J. & Dunlap, J. C. (2005) Temperature-modulated alternative splicing and promoter use in the Circadian clock gene frequency, Mol Biol Cell. 16, 5563–71.

36. Cemel, I. A., Ha, N., Schermann, G., Yonekawa, S. & Brunner, M. (2017) The coding and noncoding transcriptome of Neurospora crassa, BMC Genomics. 18, 978.

37. He, Q., Cheng, P., Yang, Y., Wang, L., Gardner, K. H. & Liu, Y. (2002) White collar-1, a DNA binding transcription factor and a light sensor, Science. 297, 840–3.

38. Smith, K. M., Sancar, G., Dekhang, R., Sullivan, C. M., Li, S., Tag, A. G., Sancar, C., Bredeweg, E. L., Priest, H. D., McCormick, R. F., Thomas, T. L., Carrington, J. C., Stajich, J. E., Bell-Pedersen, D., Brunner, M. & Freitag, M. (2010) Transcription factors in light and circadian clock signaling networks revealed by genomewide mapping of direct targets for neurospora white collar complex, Eukaryot Cell. 9, 1549–56.

39. Belden, W. J., Lewis, Z. A., Selker, E. U., Loros, J. J. & Dunlap, J. C. (2011) CHD1 remodels chromatin and influences transient DNA methylation at the clock gene frequency, PLoS Genet. 7, e1002166.

40. Mullaney, E. J., Hamer, J. E., Roberti, K. A., Yelton, M. M. & Timberlake, W. E. (1985) Primary structure of the trpC gene from Aspergillus nidulans, Mol Gen Genet. 199, 37–45.

41. Diernfellner, A. C., Schafmeier, T., Merrow, M. W. & Brunner, M. (2005) Molecular mechanism of temperature sensing by the circadian clock of Neurospora crassa, Genes Dev. 19, 1968–73.

42. Gooch, V. D., Mehra, A., Larrondo, L. F., Fox, J., Touroutoutoudis, M., Loros, J. J. & Dunlap, J. C. (2008) Fully codon-optimized luciferase uncovers novel temperature characteristics of the Neurospora clock, Eukaryot Cell. 7, 28–37.

43. Cesbron, F., Brunner, M. & Diernfellner, A. C. (2013) Light-dependent and circadian transcription dynamics in vivo recorded with a destabilized luciferase reporter in Neurospora, PLoS One. 8, e83660.

44. Bell-Pedersen, D., Shinohara, M. L., Loros, J. J. & Dunlap, J. C. (1996) Circadian clock-controlled genes isolated from Neurospora crassa are late night-to early morning-specific, Proc Natl Acad Sci U S A. 93, 13096–101.

45. Sancar, C., Sancar, G., Ha, N., Cesbron, F. & Brunner, M. (2015) Dawn-and dusk-phased circadian transcription rhythms coordinate anabolic and catabolic functions in Neurospora, BMC Biol. 13, 17.

46. Sancar, G., Sancar, C., Brugger, B., Ha, N., Sachsenheimer, T., Gin, E., Wdowik, S., Lohmann, I., Wieland, F., Hofer, T., Diernfellner, A. & Brunner, M. (2011) A global circadian repressor controls antiphasic expression of metabolic genes in Neurospora, Mol Cell. 44, 687–97.

47. Sancar, C., Ha, N., Yilmaz, R., Tesorero, R., Fisher, T., Brunner, M. & Sancar, G. (2015) Combinatorial control of light induced chromatin remodeling and gene activation in Neurospora, PLoS Genet. 11, e1005105.

48. Sancar, G., Sancar, C. & Brunner, M. (2012) Metabolic compensation of the Neurospora clock by a glucose-dependent feedback of the circadian repressor CSP1 on the core oscillator, Genes Dev. 26, 2435–42.

